# Arabidopsis ROOT UV-B SENSITIVE 1 and 2 Interact with Aminotransferases to Regulate Vitamin B6 Homeostasis

**DOI:** 10.1101/2021.03.01.433438

**Authors:** Hongyun Tong, Colin D. Leasure, Robert Yen, Xuewen Hou, Nathan O’Neil, Dylan Ting, Ying Sun, Shengwei Zhang, Yanping Tan, Elias Michael Duarte, Stacey Phan, Cinthya Ibarra, Jo-Ting Chang, Danielle Black, Tyra McCray, Nate Perry, Xinxiang Peng, Jesi Lee, Keirstinne Turcios, Anton Guliaev, Zheng-Hui He

## Abstract

Pyridoxal-5’-phosphate (PLP), the enzymatic cofactor form of Vitamin B6 (vitB6), is a versatile compound that has essential roles in metabolism. Cellular PLP homeostasic regulation is currently not well understood. Here we report that in Arabidopsis, biosynthesized PLP is sequestered by specific aminotransferases (ATs), and that the proteins ROOT UV-B SENSITIVE 1 (RUS1) and RUS2 function with ATs to regulate PLP homeostasis. The stunted growth phenotypes of *rus1* and *rus2* mutants were previously shown to be rescuable by exogenously supplied vitB6. Specific residue changes near the PLP-binding pocket in ASPARTATE AMINOTRANSFERASE2 (ASP2) also rescued *rus1* and *rus2* phenotypes. In this study, saturated suppressor screens identified 14 additional *suppressor of rus* (*sor*) alleles in four aminotransferase genes (*ASP1*, *ASP2*, *ASP3*, or *ALANIN AMINOTRANSFERASE1* (*AAT1*)), which suppressed the *rus* phenotypes to varying degrees. Each of the *sor* mutations altered an amino acid in the PLP-binding pocket of the protein, and sor proteins were found to have reduced levels of PLP conjugation. Genetic data revealed that the availability of PLP normally requires both RUS1 and RUS2, and that increasing the number of *sor* mutants additively enhanced the suppression of *rus* phenotypes. Biochemical results showed that RUS1 and RUS2 physically interacted with ATs. Our studies suggest a mechanism in which RUS1, RUS2 and specific ATs work together to regulate PLP homeostasis in Arabidopsis.

## Introduction

Vitamin B6 (vitB6) is an essential organic compound in all living cells (György, 1934; György and Eckardt, 1940; Tambasco-Studart et al., 2005). VitB6 is any of six vitamer compounds: pyridoxal (PL), pyridoxine (PN), pyridoxamine (PM), and their 5’-phosphates (PLP, PNP, PMP, respectively). PLP is the vitB6 vitamer used as a cofactor in a diverse array of enzymes. PLP-dependent enzymes make up about 4% of all of the classified enzymes by the Enzyme Commission (Mozzarelli and Bettati, 2006). These enzymes catalyze a broad range of reactions, including decarboxylation, transamination, racemization, and elimination (Percudani and Peracchi, 2003; Mooney et al., 2009). These biochemical reactions create a high demand for PLP in a wide range of physiological functions, including amino acid metabolism, glucose metabolism, lipid metabolism, and hormone synthesis (Mooney and Hellmann, 2010; Boycheva et al., 2015). Metabolic activities during early seedling establishment require a synchronized supply of vitB6 to both break down energy-storage molecules and build up new macromolecules for new cells and organs. In addition to its enzymatic functions, vitB6 has been shown to conjugate nuclear corepressor RIP410 to regulate gene expression (Huq et al., 2007), and to serve as a potent antioxidant against environmental stresses (Ehrenshaft et al., 1999; Chen and Xiong, 2005; Havaux et al., 2009; Mooney et al., 2009).

Whereas animals are unable to synthesize vitB6 themselves, plants and many other organisms directly synthesize PLP through the activity of the PLP synthase enzyme (Ehrenshaft et al., 1999; Ehrenshaft and Daub, 2001; Titiz et al., 2006; Fitzpatrick et al., 2007). In Arabidopsis the mature PLP synthase complex consists of twelve PDX1 and twelve PDX2 subunits (Leuendorf et al., 2010, 2014). PDX2 is a glutaminase that removes an ammonium from glutamine and channels it to PDX1 (Tambasco-Studart et al., 2005, 2007). PDX1 facilitates the combination of ribose-5-phosphate and glyceraldehyde-3-phosphate (or similar sugars) with the ammonium provided by PDX2 to directly create PLP as a product (Tambasco-Studart et al., 2005).

PLP can also be recovered from protein turnover and through the so-called salvage pathway (González et al., 2007). The salvage pathway involves the activity of PN/PM/PL kinase (SOS4), which phosphorylates any of the non-phosphorylated vitB6 vitamers, and PNP/PMP oxidase (PDX3), which oxidizes PNP or PMP into PLP (Shi et al., 2002; González et al., 2007; Sang et al., 2011). Additionally, PLP can be converted into PNP by the pyridoxal reductase (PLR1) enzyme (Herrero et al., 2011). All phosphorylated vitB6 vitamers can be dephosphorylated, possibly by a generic phosphatase enzyme (Huang et al., 2011). Non-PLP vitB6 vitamers are present in plant extracts at different levels, and they may originate from both external or internal sources. VitB6 vitamers released by soil microorganisms can be taken in by plant root cells, which presumably have vitB6 transporters in their plasma membranes. PMP is formed as an intermediate carrier of the amino group during transamination reactions, and the relatively high levels of PMP in plant extracts suggests that transamination reactions may only be partial in certain circumstances. A number of studies suggest that the function of the PLP salvage pathway is more than just for PLP regeneration (Shi et al., 2002; González et al., 2007; Colinas et al., 2016). The interconversions among various vitB6 vitamers through the PLP salvage pathway can provide a balance of vitB6 vitamers to meet metabolic and developmental demands. Analysis of *pdx3* mutants suggested that PMP accumulation causes an imbalance in vitB6 vitamers and affects nitrogen metabolism (Colinas et al., 2016). The PLP salvage pathway can also function to convert PLP originating from several possible sources, including the *de novo* PLP synthesis pathway and the protein turnovers of PLP-bound enzymes, to other vitB6 vitamers for storage.

The vitB6 pool may be depleted due to a combination of external and internal factors. VitB6 vitamers can be destroyed when exposed to UV-B light, and PLP is more rapidly degraded than the other vitamers (Leasure et al., 2013). As phototrophs, plants are constantly exposed to solar light, and plants must manage the PLP degradation induced by UV-B. VitB6 is a potent antioxidant compound and is depleted during the quenching of reactive oxygen species (ROS), which may accumulate under various environmental stresses. However, the major exhaustion of the vitB6 pool is the biochemical conjugation of PLP to lysine residues via a Schiff base covalent bond. This occurs in more than 100 different enzymes that are located in different cellular compartments including cytoplasm, peroxisomes, mitochondria and chloroplasts. In addition, the bioactive vitB6 vitamer, PLP, is known to pose a biochemical dilemma. While the constant supply of PLP is essential to cellular functions, PLP accumulation can cause cytotoxicity. The 4’-aldehyde of PLP can form aldimines with α-amines of amino acids and other compounds containing amino groups, ε-amino groups of lysine residues on non-B6 binding proteins, and thiazolidine adducts with sulfhydryl groups like cysteine (Ghatge et al., 2012). It is important that PLP homeostasis in a metabolically active cell is closely monitored and maintained (Xia et al., 2014). How vitB6 homeostasis is monitored and regulated during development is currently not well understood.

We have identified two Arabidopsis *<u>r</u>oot <u>u</u>v-b <u>s</u>ensitive* (*rus1* and *rus2*) mutants that are developmentally arrested post-germination (Tong et al., 2008; Leasure et al., 2009). This arrest is partially alleviated if the very-low fluence-rate UV-B present in laboratory lights is filtered out. The *rus1* and *rus2* mutant phenotypes can be more strongly rescued by increasing the levels of exogenously added vitB6, which is a standard additive to plant growth media (González et al., 2007). Further studies suggested that these two rescue conditions are related, as vitB6 is photolytically destroyed by UV-B (Leasure et al., 2011). RUS1 and RUS2 are homologous proteins that functionally work together in a complex, and they are members of a large protein family that contains DUF647 (<u>d</u>omain of <u>u</u>nknown function 647), a domain found in most eukaryotic organisms (Leasure et al., 2009). Interestingly, although *rus1* and *rus2* mutants can be partially rescued by exogenous vitB6, the endogenous total vitB6 levels, including PLP, in both mutants are comparable to those in the WT (Leasure et al., 2009), suggesting that *rus1* and *rus2* mutants may have defects in modulating vitB6 homeostasis.

The *rus1* and *rus2* mutants can serve as good genetic platforms to set up suppressor screens to look for key components involved in modulating vitB6 homeostasis. Our previous genetic analyses revealed a major player in the RUS1/RUS2-mediated PLP homeostasis pathway (Leasure et al., 2011). Mutagenesis identified several mutations in the *ASPARTATE AMINOTRANSFERASE 2* (*ASP2*) gene that behaved as second-site suppressors of the *rus* mutant phenotypes. *ASP2* encodes for cytosolic aspartate aminotransferase, a vitB6-dependent enzyme that plays a key role in nitrogen metabolism (Schultz and Coruzzi, 1995). Genetic analyses of six *asp2* mutant alleles revealed that only specific amino acid substitutions in ASP2 could suppress the *rus* mutant phenotypes. Complete knockouts of the *ASP2* gene did not show any suppression, suggesting that suppression requires the presence of the mutant asp2 proteins. Strikingly, all of the substitutions capable of suppressing *rus* mutant phenotypes reside at positions in the PLP binding pocket, suggesting that the binding of PLP to ASP2 may play a key role in vitB6 homeostasis in *Arabidopsis* (Leasure et al., 2011). (Leasure et al., 2011). These findings also imply a second function for aspartate aminotransferase, in addition to its role as an enzyme. This function appears to be highly conserved, as the expression of either *Arabidopsis* or Human cytosolic aspartate aminotransferase (also known as Human glutamate-oxaloacetate transaminase 1 or hGOT1) returned suppressed *rus1 asp2* mutant plants to a *rus1* phenotype (Leasure et al., 2011). Recently, a molecular dynamics simulation study with the hGOT1 suggested that specific mutations in the PLP binding pocket of hGOT1 result in hGOT1 dimerization misalignment and the release of PLP (Lee et al., 2018). How RUS1, RUS2 and aminotransferases function together to modulate cellular vitB6 homeostasis is the subject of this study. We took advantage of the well characterized vitB6 homeostasis-related phenotypes in *rus1* and *rus2* mutants and performed further screens to identify additional genetic suppressors. Our genetic and biochemical experiments revealed crucial roles of aminotransferases in vitB6 homeostasis. It is intriguing that aspartate aminotransferase and alanine aminotransferase, both of which are widely used as crucial clinical markers for human health, were identified as key players in vitB6 homeostasis. Our data suggest that aminotransferases serve as cellular PLP sequesters, and RUS1 and RUS2 function with these PLP sequesters to modulate PLP homeostasis.

## Results

### Extragenic Suppressors of the *rus1* and *rus2* Phenotype Are Exclusively Aminotransferases

The *RUS1* and *RUS2* genes play essential roles in *Arabidopsis* early seedling development (Leasure et al., 2009; Ge et al., 2010; Leasure et al., 2011; Yu et al., 2013). To understand how RUS1 and RUS2 are involved in early seedling development, we performed an ethyl methane sulfonate (EMS)-based mutagenesis screen to identify second-site suppressors of the *rus1* or *rus2* mutant phenotype. In this study we expanded our screen to include both the *rus1* and the *rus2* background, and we identified a total of 70 additional *<u>s</u>uppressor <u>o</u>f <u>r</u>us* (*sor*) mutants. As *rus1* and *rus2* mutants developmentally arrest post-germination, the *sor* plants in either the *rus1* or the *rus2* genetic background were identified based on their elongated roots and improved leaf production (Figure 1). All identified *sor* mutants were characterized by backcrossing them to their corresponding original *rus* mutant strain (either *rus1* or *rus2*). Allelic analyses of all 70 *sor* mutants (in the *rus1* background) were performed, which placed the *sor* mutations into 14 distinct alleles belonging to four distinct genetic complementation groups (*sor1*, *sor2*, *sor3*, and *sor4*) (Figure 1). Pairwise-cross analyses and rough mapping data suggested that the *sor* mutants were found in four different genes. We previously identified *SOR1* as *ASPARTATE AMINOTRANSFERASE2* (*ASP2*), and in addition to the four previously reported *sor1/asp2* alleles, four new *sor1/asp2* alleles were identified (Figure 1B). While six suppressor alleles were identified for *sor2* and three alleles for *sor4*, only one allele was identified for *sor3* (Figure 1C, 1D, 1E). The root lengths of 7-days-old *rus1* seedlings were less than 2% that of WT seedlings (Figure 1A). The root lengths of the *sor* mutants (in the *rus1* background) were restored to between 20-60% that of the WT (Supplemental Figure 1). Various *sor1* alleles showed different levels of developmental restoration. For example, among the eight *sor1* alleles, *sor1-4* and *sor1-7* showed stronger suppression than the other six alleles (Figure 1B). The suppression effect was also observed in shoot development. The cotyledons of the *rus1* mutant were chlorotic, but all *rus sor* double mutants displayed healthy green color and true leaf development (Figure 1).

**Figure 1.**
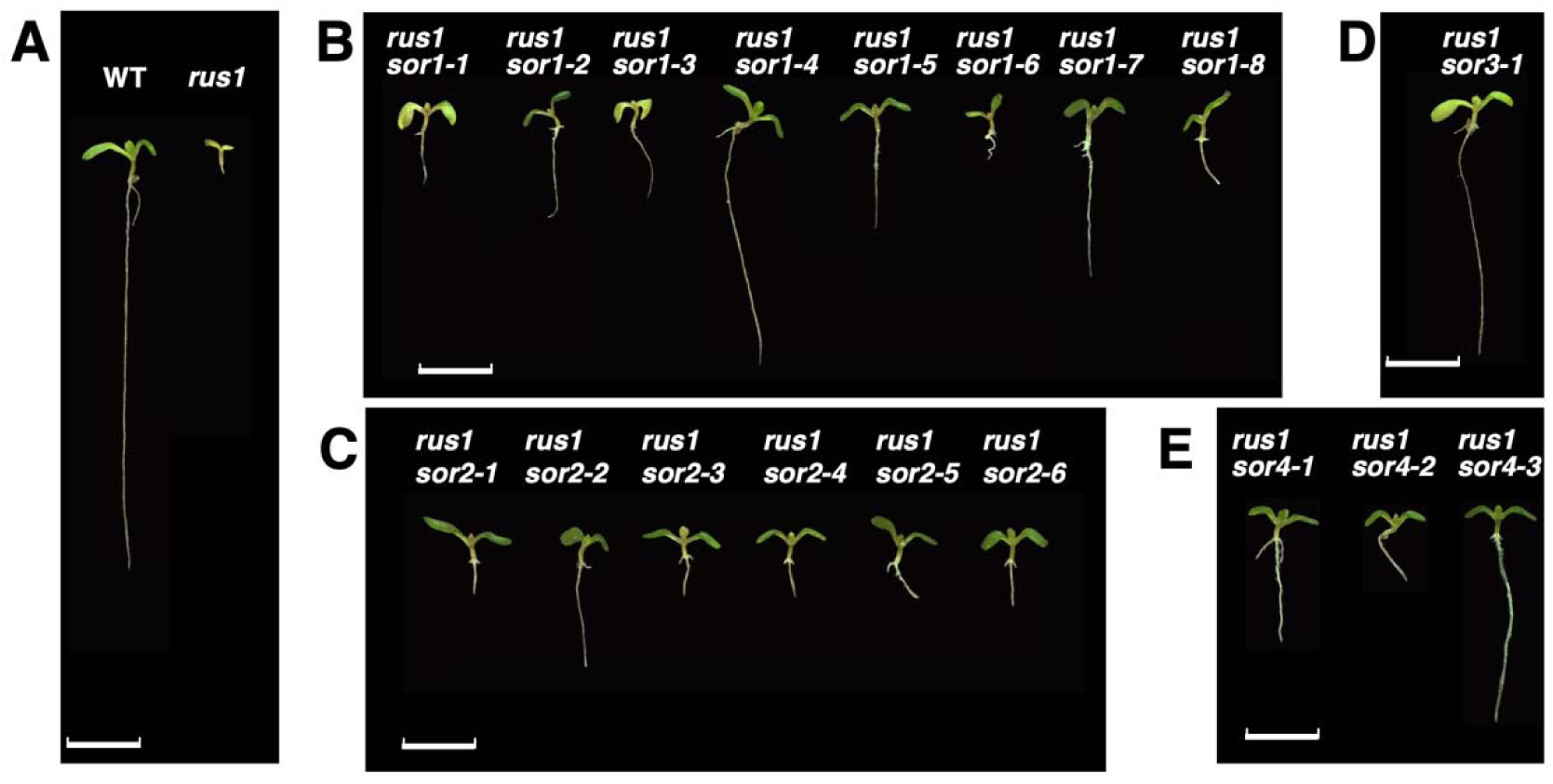
Identification of genetic suppressors of rus1. (**A**), Representative 7-day-old seedlings of WT and *rus1* (*<u>r</u>oot <u>u</u>v-b <u>s</u>ensitive1*) are shown. (**B**), Representative seedlings of eight *sor1* (*<u>s</u>uppressor-<u>o</u>f-<u>r</u>us* 1) mutant alleles in *rus1* background (allele name as labeled). (**C**), Representative seedlings of six *sor2* mutant alleles in *rus1* background (allele name as labeled). (**D**), Image of sor3-1 in *rus1* background. E. Representative seedlings of three *sor4* mutant alleles in *rus1* background (allele name as indicated). Scale bar = 5 mm.

The *sor* mutant alleles were mapped and identified based on traditional map-based cloning methods (Tong et al., 2008). Broadly spaced markers placed the four distinct genetic complementation groups onto three different chromosomes (Figure 2A). *SOR1* and *SOR3* were located on chromosome 5, whereas *SOR2* and *SOR4* were located on chromosome 2 and chromosome 1, respectively. Fine mapping and sequencing further identified all *SOR*s as genes that encode aminotransferases (Figure 2B, 2C, 2D, 2E). In addition to the previously reported *SOR1* that encodes *ASP2* (*At5g19550*), *SOR2* was identified as *ASPARTATE AMINOTRANSFERASE 1* (*ASP1*, *At2g30970*),*SOR3* as *ASPARTATE AMINOTRANSFERASE 3* (*ASP3*, *At5g11520*), and *SOR4* was identified as *ALANINE AMINOTRANSFERASE1* (*ALAAT1*, *At1g17290*) (Figure 2). A 30-kb region in the fine-mapped location was sequenced for each *sor* group, and the nature of the mutation for each *sor* allele was confirmed by sequencing. In keeping with the proper *Arabidopsis* nomenclature scheme, all *sor* alleles were renamed and renumbered as *asp1*, *asp2*, *asp3*, or *alaat1* alleles (Table 1), but “*sor”* will be used to collectively refer to *rus* suppressor mutations.

**Figure 2.**
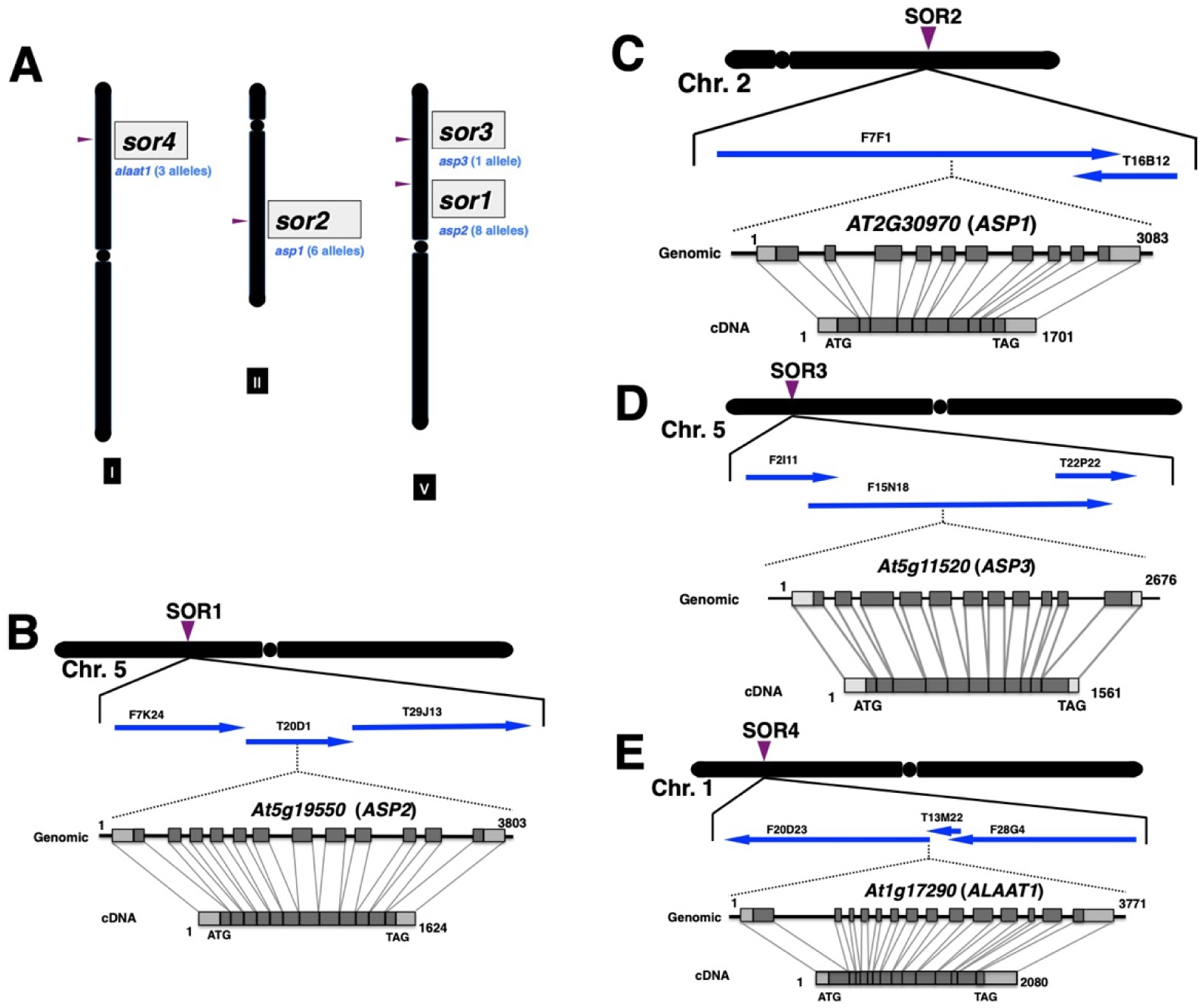
Mapping and cloning of the SOR genes. (**A**), Genomic positions of the four SOR genes were determined by genetic mapping. Arrows shows the SOR positions on their corresponding chromosomes (chromosome II and V). The number of alleles for each sor gene (sor1, sor2, sor3) and their corresponding allele names are indicated. (**B**-**E**), Map-based cloning of the SOR genes. Shaded boxes indicate exons. Position 1 is the transcriptional initiation site. Positions relative to the transcriptional initiation site in both genomic and cDNA are indicated. (**B**), The *SOR1* gene was mapped to on BAC T20D1 on chromosome 5 and was identified as At5g19550 (*ASP2*). (**C**) The *SOR2* gene was mapped to BAC F7F1 on chromosome 2 and was identified as At2g30970 (*ASP1*). (**D**), The *SOR3* gene was mapped to BAC F15N18 on chromosome 5 and was identified as At5g19550 (*ASP3*). (**E**),The *SOR4* gene was mapped to BAC T13M22 on chromosome 1 and was identified as At1g17290 (*ALAAT1*).

**Table 1:**
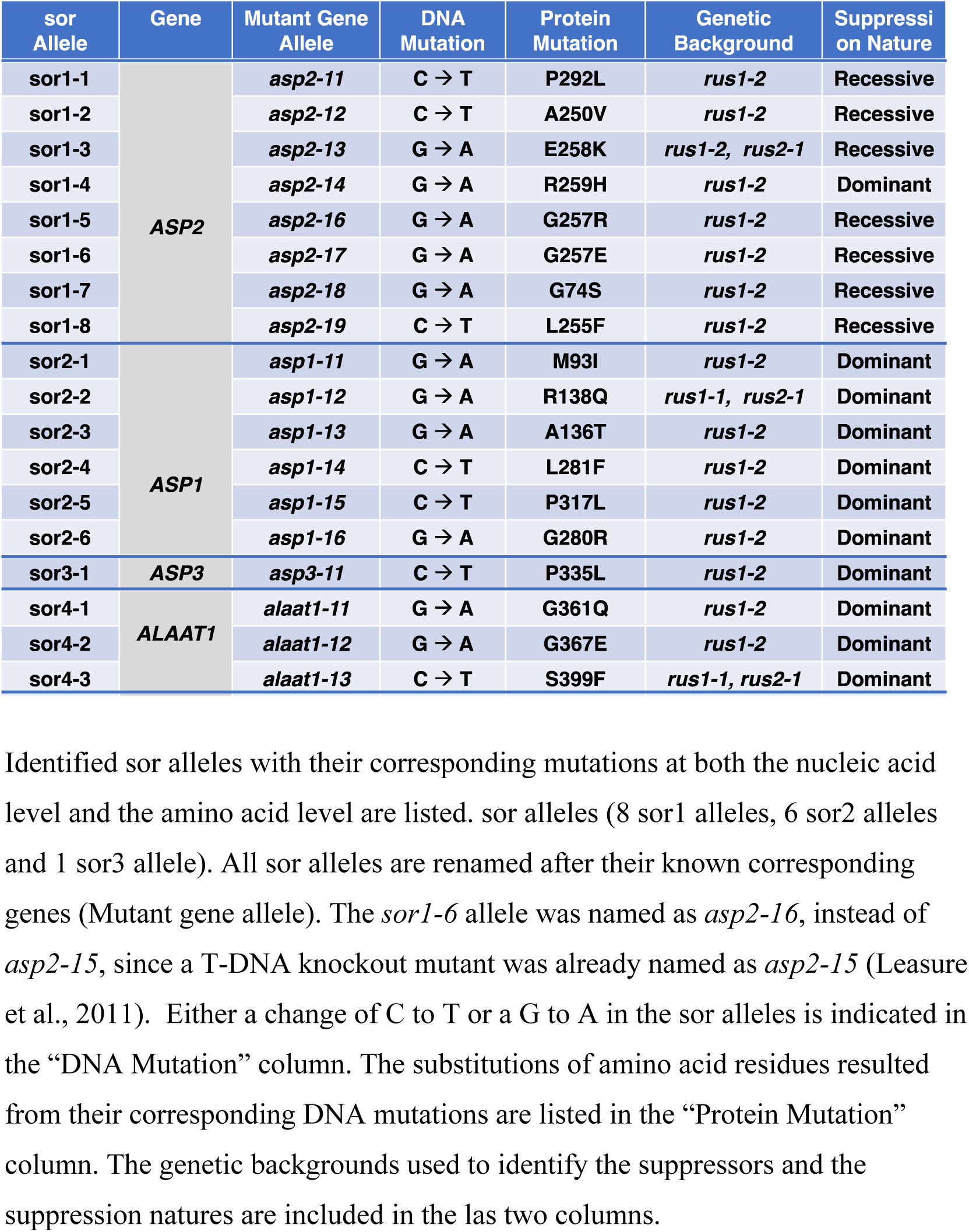
List of identified suppressors and their corresponding genes and mutations.

As expected for EMS-based mutagenesis, all detected *sor* mutations were transition mutations (either C to T or G to A), and all mutations were missense mutations in the aminotransferase genes (Table 1). *RUS1* and *RUS2* are known to work as genetic partners, and three of the four *sor* genes were identified in both the *rus1* and the *rus2* genetic background screens (Table 1), further ratifying that RUS1 and RUS2 function as partners. To generate purified *sor* alleles in either the *rus1* or the *rus2* background, all *sor* mutant alleles were backcrossed multiple times with the original mutant strain that was used to initiate the EMS-mutagenesis. Several suppressor alleles were independently identified multiple times in the various screens. For example, *asp2-13* was identified in separate genetic screens using EMS-mutagenized seeds from either the *rus1-2* or the *rus2-1* mutant background (Table 1). In both cases, the G to A mutation resulted in the codon change of GAG to CAG, and consequently a missense mutation of a Glu to a Lys. Similarly, both *asp1-12* and *alaat1-13* were independently identified in both the *rus1* and the *rus2* genetic background screens. The codon GGG that encodes G257 in ASP2 was independently hit multiple times. GGG was mutated to AGG, resulting in a G257R substitution in *asp2-16*. The same codon was mutated to GAG resulting in a G257E substitution in *asp2-17*. While only one out of the eight suppressor *asp2* alleles was dominant, all suppressor alleles for *asp1*, *asp3* and *alaat1* were dominant (Table 1). The various *rus sor* double mutants were also outcrossed to WT and subsequent progeny that carried only the *sor* mutations were identified. Under normal growth conditions, no major growth or developmental differences were observed between these *sor* single mutants and WT (Supplemental Figure 2). Our screen for *sor* mutations is likely saturated, as multiple mutations were identified in each of the *SOR* genes, with the notable exception of *ASP3.* (Table 1).

### The sor Proteins Contain Specific Amino Acid Substitutions that Impact ASP Conformation and Pyridoxal-5’-Phophate Binding

The *sor* mutations found in the *asp* genes created changes that were located primarily within two distinct regions of the proteins. One mutation hotspot region was near the PLP-binding lysine (K251 for ASP2) in the active site of the enzyme (Figure 3). In some cases, the same residue substitutions were identified multiple times. Both *asp2-19* and *asp1-14* convert the conserved leucine four amino acids after the active-site lysine into a phenylalanine (Figure 3). Another mutation hotspot was identified around forty amino acids after the active-site lysine. There are two highly-conserved prolines in this region of the aspartate aminotransferase proteins. The two prolines in this position create a kink in the protein, which is known to be essential for the proper folding of the human aspartate aminotransferase, as illustrated in its crystal structure (Figure 3B). A substitution mutation in either of the two prolines in all three ASPs (P317L for asp1-15, P292L for asp2-11, and P335L for asp3-11) resulted in *rus1* suppression. Although the primary amino acid sequence of alanine aminotransferase 1 (ALAAT1) is diverse from the aspartate aminotransferases, the amino acid changes in the three suppressor *alaat1* alleles were located in a similar pattern. Two alleles had amino acid substitutions near the active site, K369, and one allele had an amino acid substitution, S399F, in a location close to the proline site of the ASPs.

**Figure 3.**
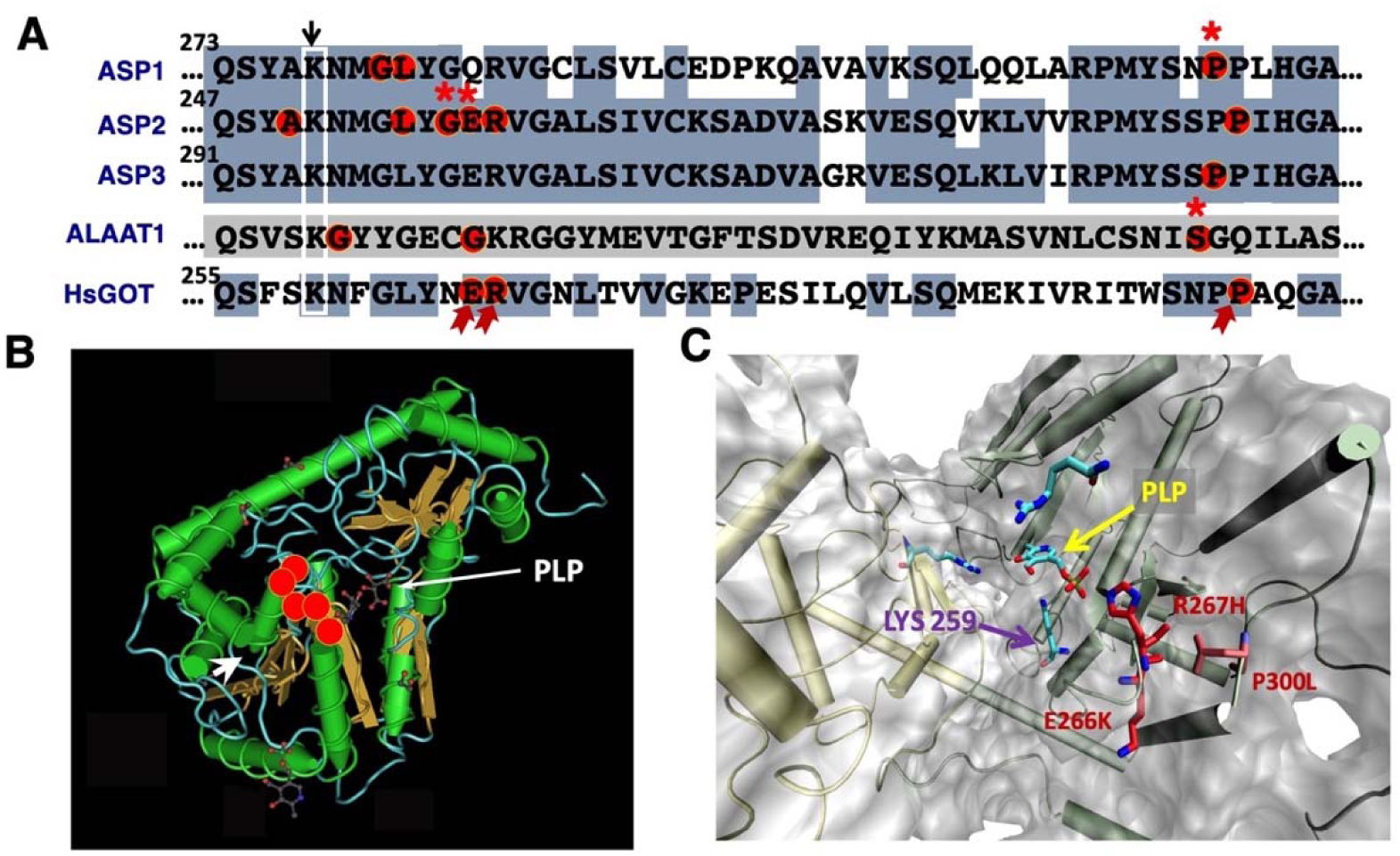
The *sor* mutants encode aminotransferases (asp1, asp2, asp3 and alaat1) with substitutions of specific amino acid residues in the vitamin B6 binding pocket. (**A**), Specific residue changes in various sor alleles. Alignment of amino acid sequences around the PLP binding site of the four aminotransferase proteins is shown. Conserved amino acid residues are shaded in light blue. Numbers indicate their corresponding amino acid positions for each aminotransferase protein. Arrow with a rectangle indicates the conserved lysine (K277 for ASP1) residue that covalently links to vitamin B via Schiff base. The substituted residues in sor mutant proteins are circled and highlighted in red circle. Multiple hot spot mutations repeatedly identified in separate suppressor screening experiments are indicated by *. (**B**), Positions of the amino acids that are substituted in the sor proteins. The conserved 3-D structure of the human aspartate aminotransferase (human glutamate-oxaloacetate transaminase, hGOT1, PDB ID 30II) is used to indicate the position of the vitamin B6 binding pocket. The circles indicate the locations of the mutated residues in the mutated sor proteins. The flat arrows are showing the beta-sheets and the round arrows for alpha helices. (**C**), Structural features of PLP binding pocket in hGOT1. WT hGOT1 has Lys259 that conjugates with PLP. Positions of both the Lys259 the PLP molecule are indicated. PLP conjugation is stabilized by key residues of E266 and R267. P300 is essential in forming the PLP binding pocket. Positions of the three substitution mutants (E266K, R267H, P300L) used in Molecular Dynamics simulation are indicated.

The specific locations and physical proximities of the substituted amino acids in the sor mutant proteins suggested that these amino acids were structurally and functionally related. The ASP enzyme works as a homodimer and each monomer has a binding pocket for PLP. Since the sor mutant proteins are aminotransferases that are known to bind PLP as a cofactor, we reasoned that the specific structural changes in the sor mutant proteins might cause PLP binding-property changes that resulted in increased PLP release from the binding pocket. A recent Molecular Dynamics (MD) simulation study with the human cytosolic aspartate aminotransferase (hGOT1) structure suggested that specific mutations within the PLP binding pocket of hGOT1 may affect hGOT1 (PDB ID 30II) dimerization and PLP binding (Lee et al., 2018). ASPs are highly conserved across all species, and the *Arabidopsis* ASP2 shares 68.8% similarity and 48.4% identity with the hGOT1 homolog (Supplemental Figure 3). We used MD to analyze three of the sor mutations using the characterized crystal structure of hGOT1 to assess whether the amino acid changes affected PLP binding. The sor mutant protein substitutions analyzed were located either within the PLP binding pocket (E266K and R267H), or at the two proline site required for correct folding of the PLP binding pocket (P300L) (Figure 3B). We hypothesized that these residues play key roles in the structural coupling between PLP and the apoprotein. To test this hypothesis, we analyzed the possible structural correlation between PLP and the PLP binding pocket throughout 1 μs MD trajectories (Figure 4). The Pearson correlation coefficient (PCC) values that represented the optimal fit between the PLP and PLP binding pocket essential for the stability of the PLP-conjugated protein were calculated. All three sor mutants simulated in hGOT1 (E266K, R267H and P300L) had much lower PCC values than WT, indicating that PLP binding at the PLP binding pocket was destabilized. While the PCC value for WT was 0.60, the PCC values were 0.24, 0.20 and 0.17 for E266K, R267H and P300L, respectively (Figure 6). The dramatically decreased PCC values for these three mutants suggested that the specific residue changes in the sor mutant proteins weakened the non-covalent bonding network supporting PLP, and disrupted the PLP-Lys binding environment causing PLP to dislodge from the active site.

**Figure 4.**
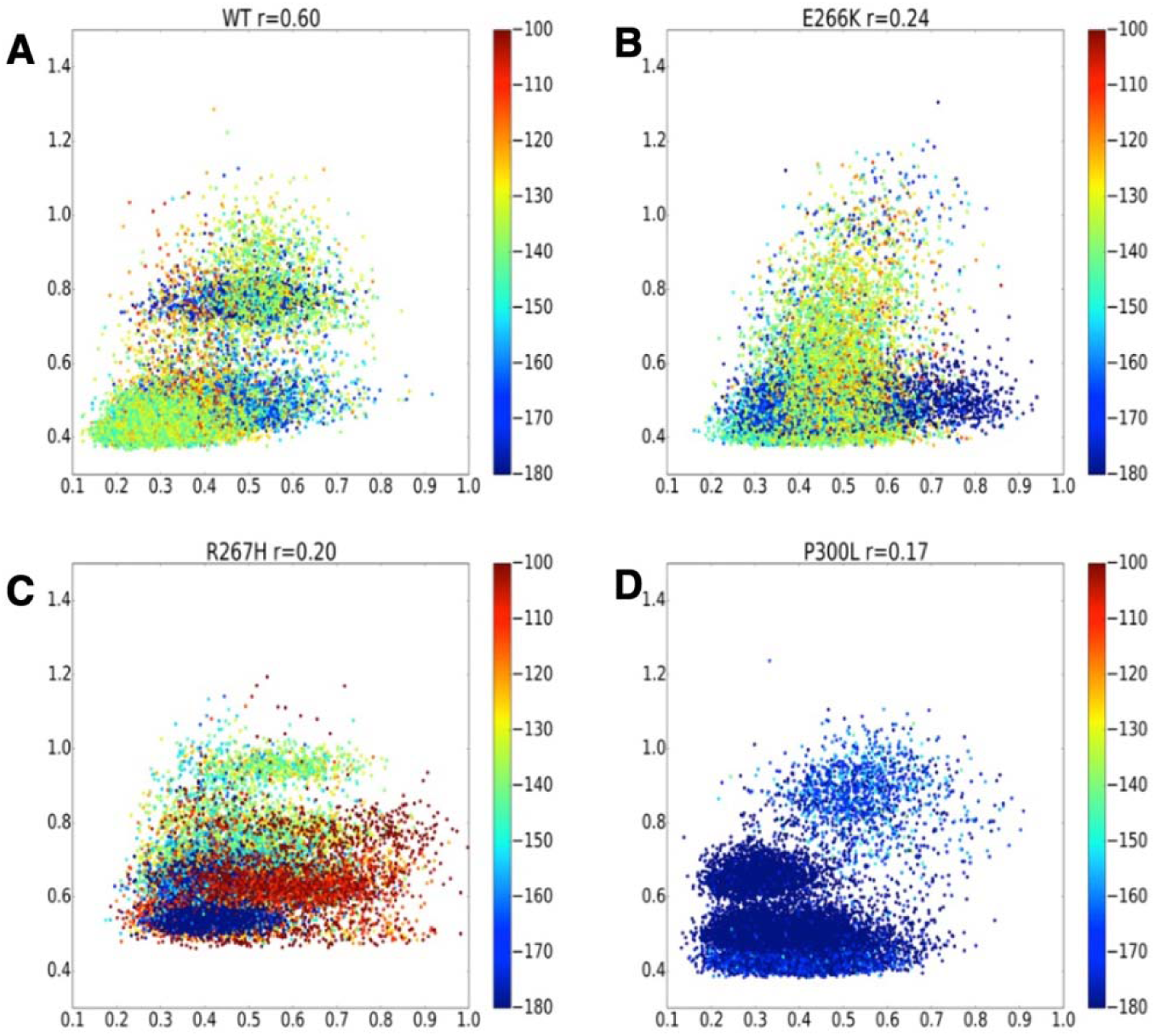
Structural correlation between PLP and PLP-binding pocket throughout 1 µs Molecular Dynamics trajectories. The structural coupling between PLP and hGOT1 was evaluated by computing Pearson correlation coefficient (PCC). Lower PPC values indicate the destabilization of the PLP at the binding pocket. Each point represents a single conformational snapshot from MD trajectory colored according to the interaction energy (left vertical color scale) between PLP and apoprotein. (**A**), PPC for the WT hGOT1 is 0.6. This value represents optimal fit between the PLP and PLP-binding pocket essential for the stability of the coenzyme and catalytic function. (**B**), PPC value for E266K mutant is 0.24. (**C**), PPC value for R267H mutant is 0.20. (**D**), PPC value for P300L mutant is 0.17. Note that the WT complex conformations are not always correspond to the lowest energy.

MD simulation data suggested that PLP binding plays critical roles in the interface between the two monomers. It is generally expected that cellular aminotransferases exist primarily as holoproteins with PLP already bound, as PLP-conjugation with an apoprotein is thermodynamically spontaneous. However, our genetic data suggested that both monomers and dimers exist, and that the dynamic binding of PLP to the monomer may serve as a mechanism for PLP sequestration. We thus hypothesized that both the monomer and the dimer co-exist *in vivo*. To test this hypothesis, immunoprecipitated aspartate aminotransferases were analyzed for their PLP conjugation status. Ten-day-old seedlings were homogenized in a protein extraction buffer containing sodium borohydrate, a compound used to reduce the Schiff base to keep PLP attached to the lysine during protein purification, digestion and subsequent peptide analysis (Billman and Diesing, 1957). Immunoprecipitated proteins were subjected to in-solution trypsin digestion and liquid chromatography/electrospray ionization-tandem mass spectrometry (LC/ESI-MS/MS) analysis. Trypsin hydrolyzes peptides at the carboxyl side following either a lysine or an arginine, unless the amino acids are followed by proline. Additionally, trypsin will not hydrolyze a peptide after a lysine if the lysine is covalently attached to PLP. Peptides generated by trypsin digestion were analyzed by LC/ESI-MS/MS. PLP-conjugated peptides were distinguished by both their longer length and the additional molecular weight gain of 231 Daltons caused by the presence of the PLP molecule itself. Peptides derived from trypsin digestion of the immunoprecipitated ASP2 protein covered more than 73% of the ASP2 protein, and two kinds of peptides containing K251 were detected (Figure 5). Peptides with the sequence TFVADGGECLIAQSYAKNMGLYGER were detected (Figure 5B), suggesting the carboxyl end of K251 was not cleaved by trypsin. PLP conjugation to K251 in these peptides was further confirmed by the additional 231 Dalton molecular weight. Peptides with the sequence TFVADGGECLIAQSYAK were also detected (Figure 5A), suggesting that both apoprotein (without PLP conjugation) and holoprotein (with PLP conjugation) co-existed *in vivo*.

**Figure 5.**
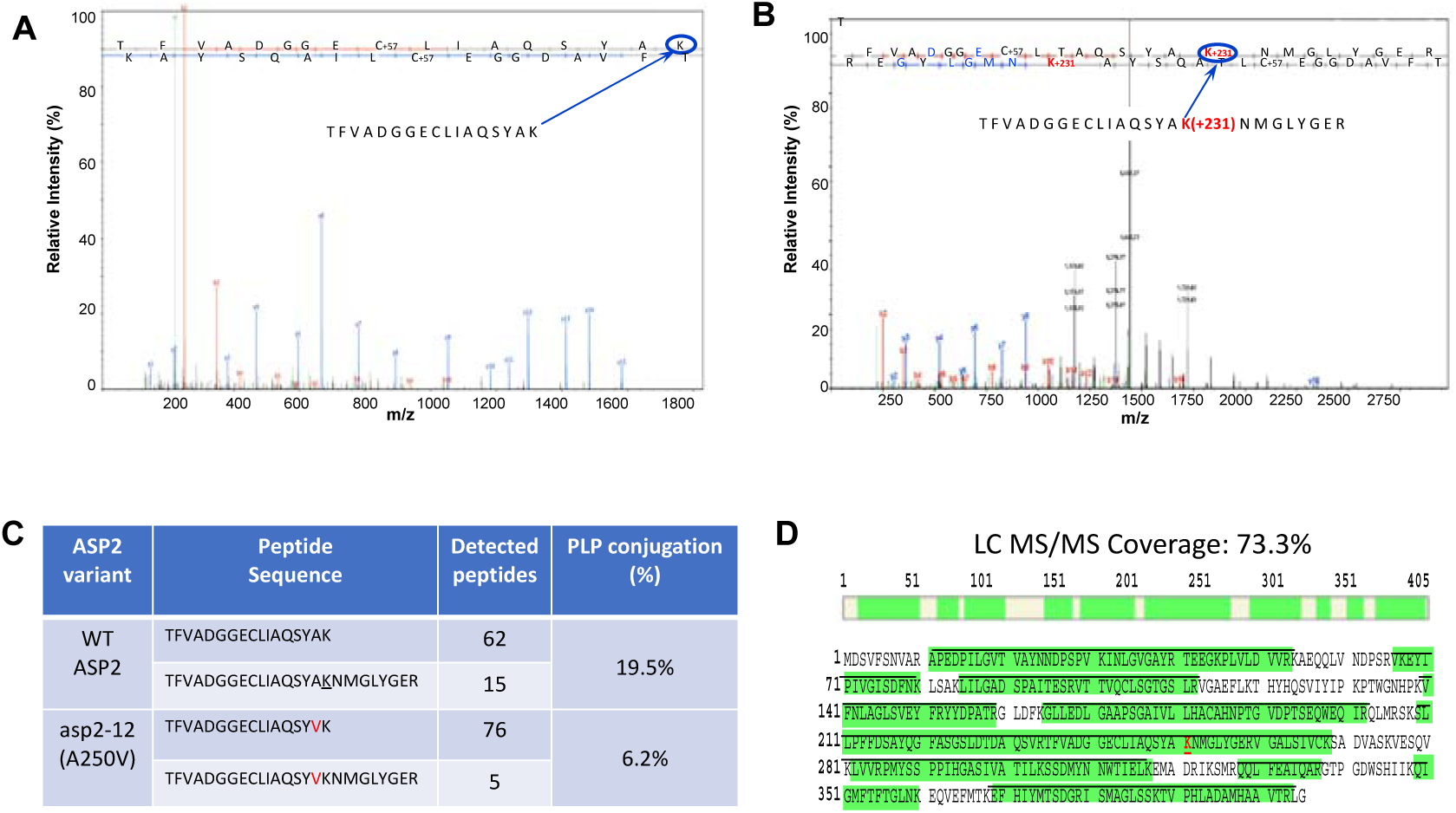
LC/ESI-MS/MS analysis of PLP conjugation in immunoprecipitated aminotransferases. Total proteins extracted from 8-day-old seedlings (either WT or sor1-2 mutant) were used for immunoprecipitation using a polyclonal anti-ASP2 antibody. (**A**), MS/MS spectrum of the expected ASP2 peptide containing K251 (TFVADGGELLIAQSYAK) is generated by trypsin digestion. The K251 site (arrow), when not covalently linked to PLP, is cleaved by trypsin. (**B**), MS/MS spectrum of the longer peptide (TFVADGGELLIAQSYAKNMGLYGER) generated when the K251 site is covalently linked to PLP and protected from trypsin digestion. The presence of PLP in this peptide gives a molecular weight gain of 231 Dalton (+231, as indicated). (**C**), PLP conjugation is affected by specific residue changes in suppressor protein (sor1-2). Numbers of peptides with or without PLP attachments detected in both WT and sor1-2 are listed and the percent of PLP conjugation is calculated (PLP conjugation ratio). (**D**), Peptide sequence of ASP2 detected by LC/ESI-MS/MS. The regions that were detected by MS/MS are highlighted. Total LC/ESI-MS/MS coverage is 73.3%. The detected sequences are indicated by underlined green letters.

Immunoprecipitated proteins from both WT and *asp2-12* seedlings were analyzed side by side. Similar to that in the WT, peptides with or without PLP attachment were detected in *asp2-12*. While 19.5% of the K251-containing peptides from the WT showed PLP modification, only 6.2% of those from sor1-2 showed PLP conjugation (Figure 5C). The PLP attachment at K251 in asp2-12 is reduced to 30% of that in the WT, suggesting that the A250V mutation caused a significant change in the PLP conjugation status. We further tested whether a similar mutation in the human aspartate aminotransferase, hGOT1, can result in similar changes in PLP conjugation. Both the WT hGOT1 and the R267H mutant were expressed in CHO cells as GFP fusion proteins that were immunoprecipitated for LC/ESI-MS/MS analysis. PLP conjugation in the R267H mutant protein was reduced to 50% of that found in WT hGOT1 protein (Supplemental Fig. 4). These results suggested that both the PLP-free monomers and the PLP-conjugated dimers co-exist for *Arabidopsis* and human aspartate aminotransferases, and that the PLP-free monomers are the primary forms for both the WT and the mutant proteins.

### Additive Suppression Effects of Suppressors on *rus* Mutants

The reduced PLP binding in suppressor asp proteins is likely to be the underlying mechanism for *rus* suppression, as a total knockout of ASP2 did not suppress *rus* mutants (Leasure et al., 2011). We hypothesized that, in the absence of functional RUS1 or RUS2 proteins, the structural changes in the PLP binding pocket of the suppressor asp proteins allow more PLP to be released from the binding pocket. To test this hypothesis, we performed genetic crosses to create a *rus1* mutant that carried more than one *sor* mutation. Multiple crosses were performed and specific genetic markers were used to isolate various homozygous double, triple and quadruple mutants (Figure 6). Suppression of the *rus1* developmental arrest phenotype was dramatically enhanced in triple and quadruple mutants. Triple mutants (either *rus1 asp2 asp1* or *rus1 asp2 alaat1*) or quadruple mutant (*rus1 asp2 asp1 alaat1*) almost completely restored the *rus1* mutant to normal seedling development that was comparable to WT. There were no apparent differences between triple or quadruple mutants and WT when both true leaf formation and chloroplast development were examined. The root lengths in of the triple and quadruple mutants were also fully restored to that of the WT (Figure 5B). The additive effects of the *sor* mutants suggested that each sor protein may be able to independently provide additional vitB6 in the *rus* mutant. This additive phenotypic restoration was reminiscent of how *rus1* can be rescued by exogenously added vitB6 in a dosage-dependent manner (Leasure et al., 2011).

**Figure 6.**
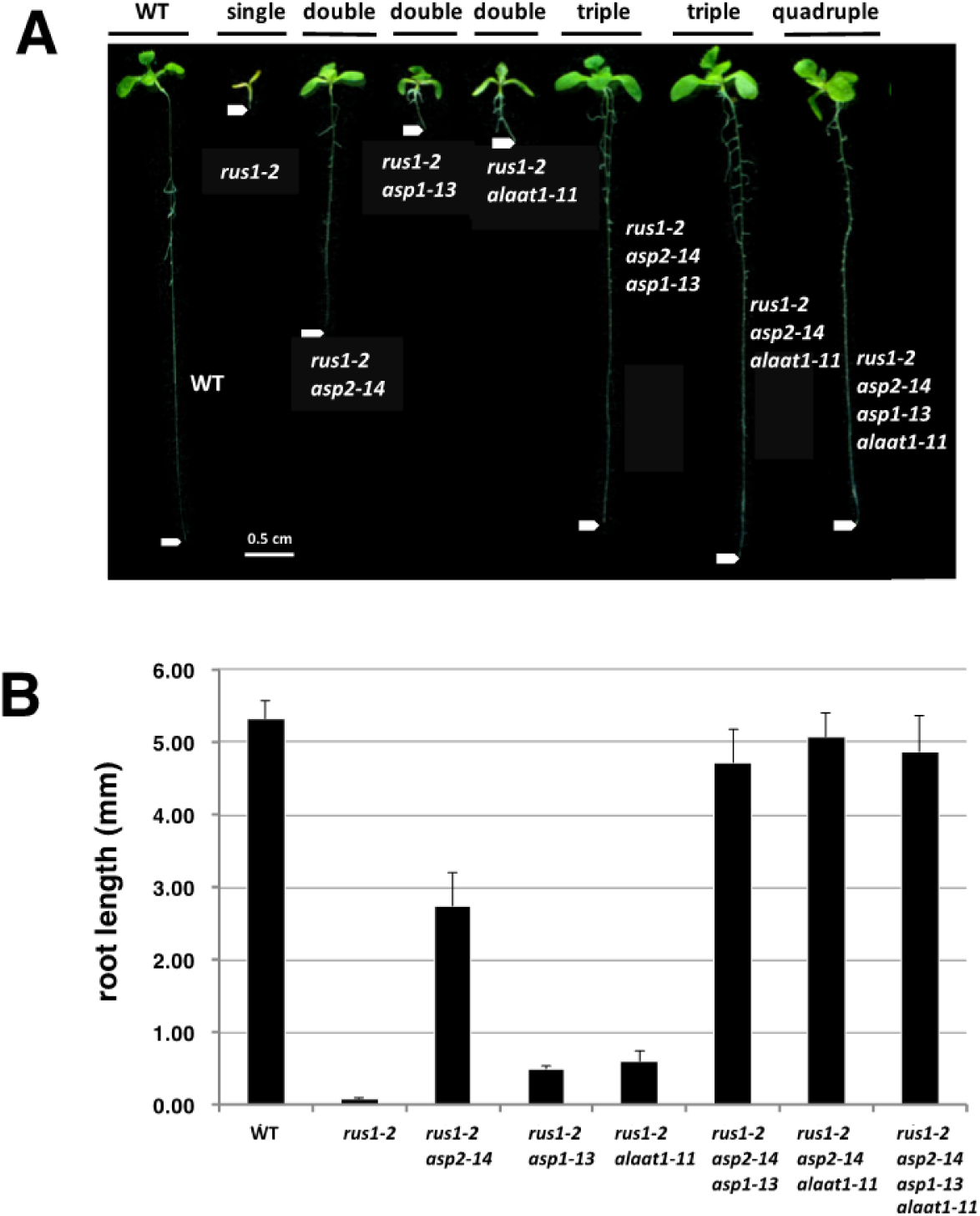
Synergistic genetic interactions among sor1,sor2, and sor4. (**A**), Representative seedlings of 9-day-old WT (WT), *rus1* (*rus1*), *rus1 sor1-1* double mutant, *rus1 sor2-1* double mutant, *rus1 sor1-1 sor2-1* triple mutant, and *rus1 sor1-1 sor2-1 sor4-1* quadruple mutant are shown (as labeled). Arrowheads indicate the positions of the root tips. Scale bar = 5 mm. (**B**), Measurements of tap root lengths in WT and the various mutants as indicated. Error bars = SE

### Role of RUS1 and RUS2 for PLP Retrieval from Aminotransferases

The *rus1* and *rus2* mutants can be rescued by externally added vitB6, although the steady state levels of vitB6 were compatible to those in the WT (Leasure et al., 2011). Our LC/ESI-MS/MS assays suggested that the suppressor proteins carried less bound PLP. These results suggested that RUS1 and RUS2 play specific roles in releasing PLP from the aminotransferases to regulate vitB6 homeostasis. It is possible that the *rus1* and *rus2* mutants exhibit a vitB6-deficency phenotype because of limited release of PLP from aminotransferases. We hypothesized that the survival of *rus1* or *rus2* mutants depends on a minimum level of PLP supplied by the vitB6 salvage pathway. To test this hypothesis, we crossed *rus1* or *rus2* with mutants that lack one of the key vitB6 salvage pathway players: SOS4 and PDX3. Both *SOS4* and *PDX3* encode enzymes that are essential for converting the various B6 vitamers to the bioactive pyridoxal 5’-phosphate (PLP) (Figure 7G). *sos4* or *pdx3* mutants show subtle phenotypes, but are still able to complete reproduction as PLP can be directly supplied by the *de novo* PLP biosynthesis pathway. The *de novo* pathway creates PLP as a product, and involves the function of the PLP synthase enzyme that consists of twelve PDX1 and PDX2 proteins each (Leuendorf et al., 2014). Homozygous *pdx2* knockout mutants were shown to be embryon lethal, which strongly suggests that plants cannot survive without PLP synthase activity. The *pdx2* mutation can only be maintained through a heterozygous *pdx2* mutant (*pdx2*/+) (Tambasco-Studart et al., 2005). Approximately 25% of embryos in young developing siliques of *pdx2*/+ plants failed to develop, which led to a white ovule phenotype (Figure 7B). We observed that homozygous *rus1-2*, *sos4-1*, and *pdx3-2* single mutants all had normal embryonic development (Figure 7A, 7C, 7E). We attempted to create homozygous *rus1-2 sos4-1* or *rus1-2 pdx3-2* double mutants through multiple genetic cross attempts. No homozygous double mutants could be identified in the segregating progeny of the selfed plants with a genotype of either *rus1-2 sos4-1*/+ or *rus1-2* pdx3-2/+ (*i.e.*, homozygous for the *rus1-2* mutation and heterozygous for either the *sos4-1* or the *pdx3-2* mutation). We suspected that homozygous double mutants for either *rus1-2 sos4-1* or *rus1-2 pdx3-2* were embryo lethal, similar to the homozygous *pdx2* mutant. To test this possibility, we analyzed embryonic development in plants with a genotype of either *rus1-2 sos4-1*/+ or *rus1-2 pdx3-2*/+. About 25% of embryos failed to develop in the siliques of both the *rus1-2 sos4-1*/+ and the *rus1-2 pdx3-2*/+ plants (Figure 7D, F). These results suggested that homozygous double mutants of either *rus1-2 sos4-1* or *rus1-2 pdx3-2* were embryo lethal.

**Figure 7.**
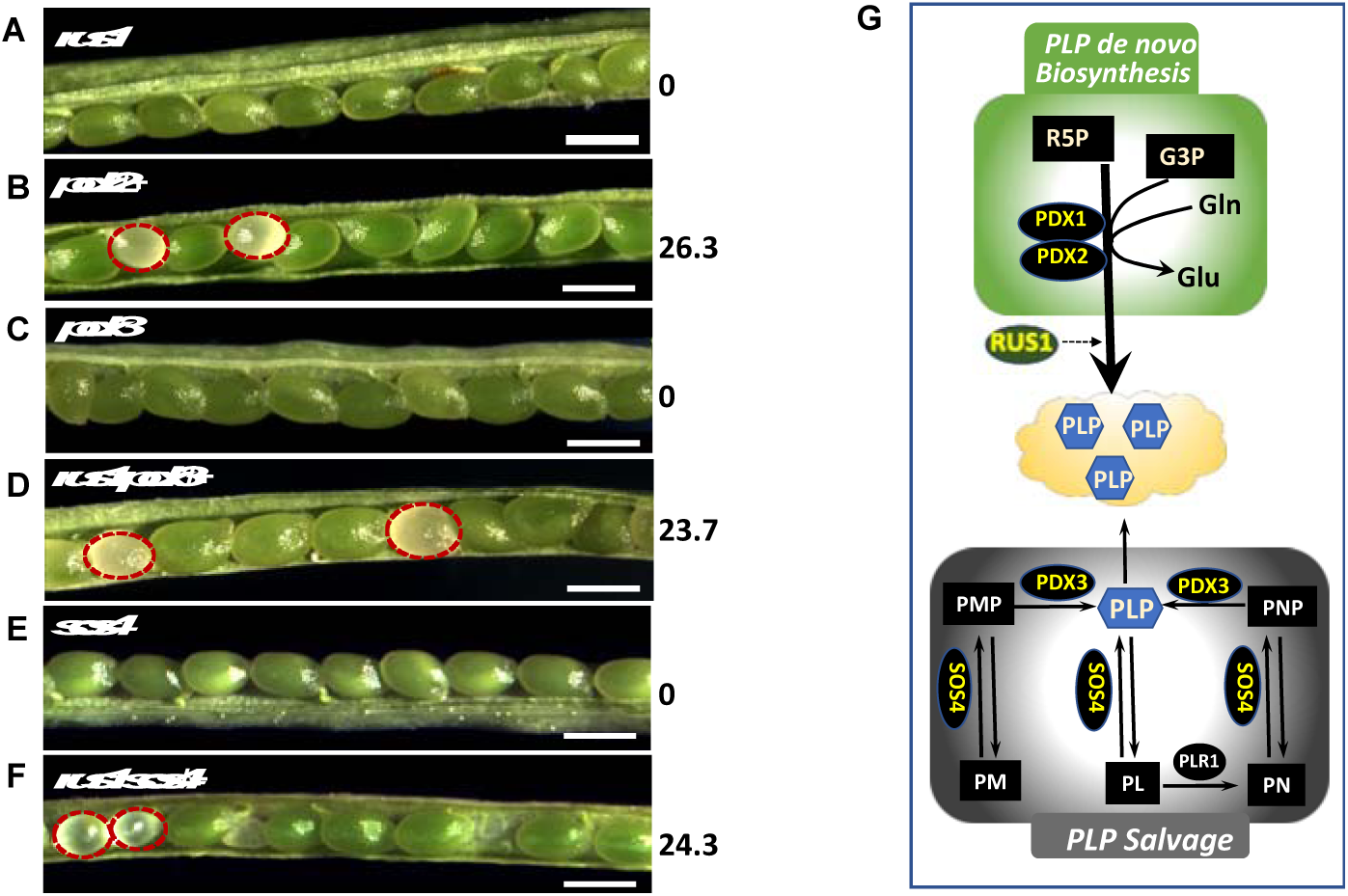
RUS1 is essential for embryonic development in mutants defective in B6 vitamer interconversions. (**A-F**), Images of embryos in opened siliques. Albino embryo numbers are scored and the percent of albino embryo is listed on the right column (percent of albino embryo). Bar = 0.5 mm. (**A**), Embryos in silique of a plant homozygous for *rus1* (*rus1*). (**B**), Embryos in silique of a plant heterozygous for *pdx2* (*pdx2*/+). Embryos homozygous for *pdx2* show albino phenotype (highlighted by a circle). (**C**), Embryos in silique of a plant homozygous for *pdx3* (*pdx3*). (**D**), Embryos in silique of a plant homozygous for *rus1* and heterozygous for *pdx3* (*rus1 pdx3*/+). Embryos homozygous for both *rus1* and *pdx3* show albino phenotype (highlighted by a circle). (**E**), Embryos in silique of a plant homozygous for *sos4* (*sos4*). (**F**), Embryos in silique of a plant homozygous for *rus1* and heterozygous for *sos4* (*rus1 sos4*/+). Embryos homozygous for both *rus1* and *sos4* show albino phenotype (highlighted by a circle). (**G**), Diagram illustrating both the de novo PLP biosynthesis pathway and the PLP salvage pathway. PLP can be directly synthesized from ribose 5-phosphate (R5P), glyceraldehyde 3-phosphate (G3P), and glutamine (Glu) via the PLP synthase complex made of PDX1 and PDX2. PLP can also be regenerated by the salvage pathway, components of which include PLR1 (pyridoxal reductase), SOS4 (PN/PM/PL kinase), and PDX3 (PNP/PMP oxidase). Genetic interaction data place RUS1at downstream of the PLP de novo biosynthesis pathway (dotted line arrow).

### RUS1 and RUS2 Interactions with Aspartate Aminotransferases

RUS1 physically interacts with RUS2 and the interactions are required for their functions in early seedling development (Leasure et al., 2009). Since RUS1 and RUS2 showed strong genetic interactions with ASPs, we hypothesized that RUS1 and RUS2 have direct protein-protein interactions with ASPs. To test this hypothesis, we performed yeast two-hybrid assays to investigate the possible interactions of either RUS1 or RUS2 with ASP2. The suppressor protein, asp2-11, was also included to test whether the specific residue change in asp2-11 would affect the interactions. RUS1 is known to strongly interact with RUS2 (Leasure et al., 2011), and this interaction was included as a positive control in the assay (Figure 8A). ASP2 works as a homodimer, and our assay confirmed that ASP2 indeed interacted with itself very strongly (Figure 8A). The single amino acid residue substitution in asp2-11 significantly weakened the direct interaction between the WT ASP2 and asp2-11 protein. Little interactions were detected for asp2-11 itself, suggesting that homodimer formation was essentially abolished by the single residue substitution in asp2-11. Interestingly, RUS1 interacted with both ASP2 and asp2-11, with a slightly stronger interaction with asp2-11. While RUS2 did not interact with ASP2, it interacted with the asp2-11 protein.

**Figure 8.**
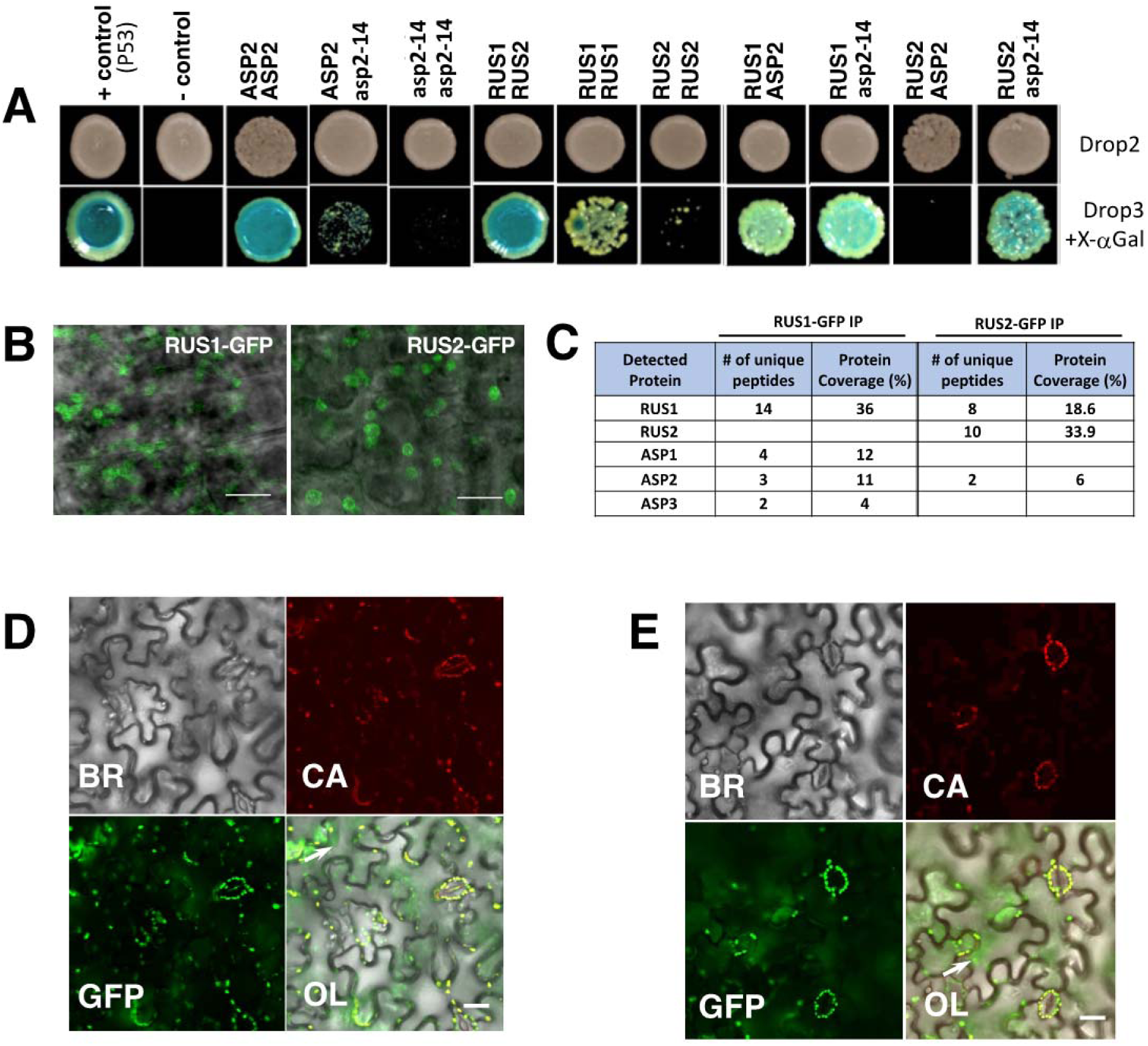
RUS1 and RUS2 interact with each other and with aspartate amino transferases. (**A**), Images of yeast two-hybrid assays. Images of spotted yeast colonies containing various paired AD (left) and BD (right) combinations on the Drop2 media and the Drop3 media containing 5-Bromo-4-chloro-3-indolyl-a-D-galactoside (X-Gal) are shown. AD-RecT was used with either the BD-p53 (+Control) or the BD-Lam (-Control) for positive and negative controls, respectively. **(B)**, Confocal localization of RUS1-GFP and RUS2-GFP as indicated. Overlay images of GFP on brightfield are shown. Scale bar = 20 nm. (**C**), Analysis of *in viv*o protein interactions by co-immunoprecipitations. Proteins immunoprecipitated by a Nanotrap GFP antibody from both the RUS1-GFP and the RUS2-GFP seedlings were analyzed by LC/ESI-MS/MS. Detected proteins with their coverages (protein coverage %) and unique peptides (# of unique peptides) are listed. **(D)** and **(E)**, Protein-protein interactions visualized by bimolecular fluorescence complementation (BiFC) assays in tobacco (*N. benthamiana*) epidermal leaf cells. GFP fluorescence images of tobacco leaf epidermal cells co-transformed either with the CCFP-RUS1 and NYFP-RUS2 constructs (D) or with the CCFP-RUS1 and NYFP-ASP2 constructs (E) are shown. Arrows indicate representative BiFC signals. BR: brightfield. CA: chlorophyll-autofluorescence. GFP: BiFC detection OL: overlay. Scale bar = 50 nm.

To verify the protein-protein interactions shown in the yeast two-hybrid assays, transgenic plants carrying either RUS1-GFP or RUS2-GFP were used for immunoprecipitation experiments. RUS1-GFP and RUS2-GFP were used to complement the *rus1* and *rus2* mutants, respectively (Tong et al., 2008; Leasure et al., 2009). High-resolution Confocal (ZEISS LSM 980 with Airyscan 2) analysis showed that RUS1-GFP and RUS2-GFP share similar localization patterns in root cells of the transgenic plants (Fig. 8B). RUS1 and RUS2 showed The immunoprecipitated proteins pulled down by an anti-GFP antibody were trypsin-digested, and the resulting peptides were analyzed by LC/ESI-MS/MS. A number of specific peptides for both RUS1 and RUS2 were detected (Figure 8C). There were 14 and 8 unique peptides identified for RUS1 and RUS2, respectively. RUS1 was detected in the immunoprecipitated samples from both RUS1-GFP and RUS2-GFP plants suggesting that RUS1 and RUS2 interacted *in vivo*. Interestingly, the three ASPs in this study (ASP1, ASP2, and ASP3) were detected in the RUS1-GFP immunoprecipitated samples. ASP2 was also detected in the RUS2-GFP immunoprecipitated sample. These results suggested that RUS1, RUS2 and ASPs have physical interactions *in vivo*. Bimolecular fluorescence complementation (BiFC) assays were used to directly test the in vivo interactions between RUS1 and RUS2, and between RUS1 and ASP2. Cells co-expressing RUS1 fused to the C-terminal half of CFP (CCFP-RUS1) and RUS2 fused to the N-terminal half of YFP (NYFP-RUS2) showed strong fluorescence signals (Fig. 8D). Cells co-expressing RUS1 fused to the C-terminal half of CFP (CCFP-RUS1) and ASP2 fused to the N-terminal half of YFP (NYFP-ASP2) also showed strong fluorescence signals (Fig. 8E). Taken together, results from the yeast two-hybrid assays, the immunoprecipitation experiment, and the BiFC assays demonstrated physical interactions between RUS1 and RUS2, and RUS1 and ASP2.

## Discussion

This study reports on a mechanism in which RUS1, RUS2 and certain aminotransferases function together to maintain vitB6 homeostasis in *Arabidopsis*. The vitB6-dependent phenotypes of *rus1* and *rus2* provided an effective platform to study vitB6 homeostasis. Our screens for genetic suppressors of the *rus1* and *rus2* mutants specifically identified three aspartate aminotransferases and one alanine aminotransferases as key players in vitB6 homeostasis. PLP, the B6 vitamer that functions as an enzymatic cofactor, is essential for the function of more than 100 enzymes, and B6 vitamers are also potent antioxidants important for photoprotection and stress responses. Despite the importance of vitB6, excessive levels of free cellular PLP can be damaging to cells, as PLP can form non-specific covalent bonds to lysines and sulfhydryl groups. The results reported in this study suggest a mechanism for how plants solved this biochemical dilemma.

Our experimental evidence suggests that PLP synthesized by the *de novo* pathway is sequestered by the aminotransferases, and that subsequent PLP release requires both RUS1 and RUS2. First, the *rus1* and *rus2* mutants behave like vitB6-deficient mutants despite having endogenous levels of vitB6 vitamers, including PLP, that are comparable to those in the WT (Leasure et al., 2011). Second, our saturated genetic screens independently and repeatedly identified specific aminotransferases as *suppressor of rus* (*sor*) gene, and all *sor* mutants created amino acid changes that affected the PLP binding pocket of the enzyme. Third, the combination of multiple *sor* genes resulted in additive genetic suppression of the *rus* phenotype, which suggests that the changes in the PLP binding pockets of the various *rus* suppressors each made some quantity of PLP available. Fourth, both *rus1 pdx3* and *rus1 sos4* double mutants were embryo lethal, which was the same phenotype seen in *pdx2* mutants that lack the *de novo* synthesis pathway. Fifth, LC/ESI-MS/MS analyses of immunoprecipitated ASP proteins found that a large portion of the aminotransferase proteins existed as the PLP-lacking apoprotein, and that a larger proportion of the suppressor proteins lacked PLP when compared to that of the WT proteins. Lastly, immunoprecipitation experiments showed that RUS1 and aspartate aminotransferase physically interacted *in vivo*. Overall, our results strongly suggest that specific aminotransferases have dual functions - in addition to their roles as enzymes, specific aminotransferases may function as PLP sequesters.

### Mutations in the aminotransferases suppress *rus* phenotypes

We performed screens for second-site suppressors of *rus1* or *rus2* mutant phenotypes, which identified suppressor mutations in four different genes. Interestingly, all *sor* mutants were determined to be in either *ASPARTATE AMINOTRANSFERASE1* (*ASP1*), *ASP2*, *ASP3*, or *ALANINE AMINOTRANSFERASE1* (*ALAAT1*). Analysis of the *sor* mutations revealed they all created residue substitutions in the PLP binding pocket of the enzyme. Our previous study of *rus* suppressor mutants in the *asp2* gene revealed that a mutation outside of the PLP binding pocket, and even a complete knock-out allele of *asp2*, each failed to suppress the *rus* phenotypes (Leasure et al., 2011). Taken together, our current and past results demonstrated that *rus* suppression requires changes in the PLP binding pocket of an aminotransferase enzymes, and is not a consequence of reduced aminotransferase activity or abundance.

The conjugation of PLP to a lysine in the PLP binding pocket via a Schiff base requires proper structural conformation in the apoprotein. The amino acid sequences in the regions responsible for forming the PLP binding pocket of aminotransferase enzymes are highly conserved, with more than 92% identity found among the aspartate aminotransferases analyzed (Supplemental Figure 3). Our Molecular Dynamics simulation studies using the known crystal structure of hGOT1 suggested that the amino acid substitutions we tested in the PLP binding pocket reduced the ability of the enzyme to bind to PLP. Consistent with this computer-based prediction, our LC/ESI-MS/MS data found that a smaller proportion of a suppressor asp2 protein had bound PLP, as compared to the WT. Furthermore, triple (*rus1 asp2 asp1* or *rus1 asp2 alaat1*) and quadruple mutant (*rus1 asp2 asp1 alaat1*) plants had increasingly restored growth, as compared to the double mutant (*rus1 asp2*; *rus1 asp1*; *rus1 alaat1*) plants. These growth differences suggested that in the absence of functional RUS1 and RUS2, each sor protein may be able to independently contribute more available PLP.

### Aminotransferase isoforms

Enzymatically functional ATs are homodimers with each monomer binding to one PLP. The binding of PLP to the apoprotein monomers is required for the subsequent formation of the holoenzyme dimers. PLP association with an AT monomer induces protein conformational change in aminotransferases and creates the required interface for the two monomers to bind Lee et al., 2018). Aminotransferases are ancient enzymes that are highly conserved across all life kingdoms. For example, the *Arabidopsis* ASP2 protein shares more than 48% amino acid sequence identity with the human aspartate aminotransferase known as hGOT1 (human glutamic-oxaloacetic transaminase 1) (Supplemental Figure 3). There are five members of the Arabidopsis *ASP* gene family (Miesak and Coruzzi, 2002). Surprisingly, no obvious phenotypes were observed in any *asp* mutant, except for some moderate growth retardation phenotypes in certain *asp2* mutants (Miesak and Coruzzi, 2002). It is possible that the ASP isoforms have redundant roles in catalyzing transamination reactions. Our results also showed that *asp* mutants that lacked aminotransferase activity were phenotypically indistinguishable from the WT (Supplemental Figure 2).

Recent protein localization data suggested that the ASP isoforms may not be confined to a single cellular compartment, as was previously proposed. For example, proteomic analyses reported that ASP2, (which was previously reported as being confined to the cytosol), was localized to the cell walls, plasmodesmata, and various membranes including the plasma membrane and the vacuolar membrane (Hooper et al., 2017). Besides the expected localization of ASP3 to the chloroplast, proteomic analyses found ASP3 to be additionally localized to mitochondria, peroxisomes, and membranes. Alanine aminotransferase 1 (ALAAT1) is known to be localized to both mitochondria and chloroplasts (Carrie et al., 2009). ASP1, ASP2 and ASP3 are highly expressed in young developing seedlings. Our histological GUS assays showed that while ASP2 is ubiquitously expressed in young developing seedlings, the expressions of both ASP1 and ASP3 were more localized to cotyledons and hypocotyls (Supplemental Figure 5). The overlapping and specific expressions of the three analyzed ASPs potentially explained why the loss of one ASP member can be enzymatically compensated by other members.

### RUS1 and RUS2 physically interact with aminotransferases *in vivo*

RUS1 and RUS2 were shown by yeast two-hybrid assays to interact with each other, and their interaction is required for their *in vivo* functions (Leasure et al., 2009). Our *in vivo* immunoprecipitation results further confirmed that RUS1 and RUS2 physically interact. RUS1 was positively identified when RUS2 was used as the bait in immunoprecipitation. RUS2 was not detected in the samples immunoprecipitated when RUS1 was used as the bait. All genetic and biochemical evidence strongly supports the conclusion that RUS1 and RUS2 work as partners in regulating PLP homeostasis. Our yeast two-hybrid assays showed that RUS1 interacts with both the WT ASP2 and asp2-11 (Figure 7). It is thus highly likely that RUS1 physically interacts with both the B6-apo monomer and the B6-bound holoenzyme dimer. RUS1 immunoprecipitation experiments using RUS1 as the bait pulled down ASP1, ASP2 and ASP3 (Figure 7). PLP conjugation assays showed that both PLP-conjugated and PLP-free aminotransferase proteins were detected in the precipitated samples. Interestingly, RUS2 interacted with asp2-11but not with ASP2 and ASP2 was pulled down when RUS2 was used as the bait. These interactive results suggest whereas RUS1 directly and physically interacts with aminotransferases regardless their PLP conjugation status, RUS2 only physically interacts with ASP proteins in their B6-apo monomer forms. The association of RUS2 with ASP2, as shown in the immunoprecipitation experiment, must thus be indirectly through RUS1. It is interesting to note that both RUS1 and RUS2 may have differential binding affinities with aminotransferases depending on the PLP conjugation status in these enzymes. A multi-protein complex containing these players is possible as RUS1 forms dimers or multimers with RUS2 and aminotransferases are known to work as dimers. It is possible that RUS1 and RUS2 sense PLP homeostasis by differentially binding to aminotransferases in their monomer (PLP-free) or dimer (PLP-bound) forms. Further studies will be needed to understand the exact makeup and the stoichiometry of such a protein complex.

### Production of PLP in plants

PLP is the product of the reaction carried out by the PLP synthase enzyme, which functions in the *de novo* synthesis of PLP. PLP synthase is made of twelve each PDX1 and PDX2 proteins, which combine ribose-5-phophate and glyceraldehyde-3-phosphate with an ammonium group taken from glutamine to produce PLP. *De novo* synthesis of PLP is essential in *Arabidopsis*, as *pdx2* mutants are embryo lethal, which suggests that the plant cannot live without a functioning PLP synthase enzyme. The majority of cellular PLP originates from self-synthesis within the plant, although B6 vitamers can also be scavenged from the soil. The vitB6 salvage pathway includes all of the enzymes required to convert the various B6 vitamers into each other, and it can also form PLP needed for cellular functions. PN/PM/PL kinase, PNP/PMP oxidase, pyridoxal reductase, aminotransferases, and phosphatases are used in the salvage pathway to interconvert the B6 vitamers. Despite its name, the salvage pathway can also convert PLP into the other forms of vitB6, presumably for safer storage and/or use in other cellular needs. For example, pyridoxal reductase can convert PLP (or PL) into PNP (or PN), and PLP is converted into PMP by the side activities of PLP-dependent enzymes (Mason et al., 1969; Gerdes et al., 2012). Studies in *E. coli* found that pyridoxal kinase can bind to and subsequently transfer PLP to an enzyme apoprotein (Ghatge et al., 2012).

Our data strongly suggest that RUS1 and RUS2 are essential in making aminotransferase-sequestered PLP available to the cell. The fact that the *rus1* and *rus2* mutants can survive at all suggests that a limited amount of PLP can be made available by the salvage pathway. *rus1 pdx3* and *rus1 sos4* mutants were embryo lethal, suggesting that when RUS1/RUS2 complex is unable to retrieve PLP from aminotransferases, the salvage pathway become essential. It is likely then that while much of the newly synthesized PLP is sequestered by aminotransferases, some quantity is funneled into the salvage pathway and converted into other B6 vitamers. The plants cannot overcome the loss of both the retrieval of PLP from aminotransferases and the recovery of PLP from the salvage pathway, leading to embryo lethality.

### Second role for aminotransferases in PLP sequestration

*rus1* and *rus2* mutants have a strong vitB6-deficient phenotype, and partially recover when exogenous vitB6 levels are increased in the growth media. The B6 vitamer levels were found to be remarkably similar to those in the WT (Leasure et al., 2011). which controverted the obvious explanation that the mutants had reduced vitB6. Since both the *de novo* PLP synthesis pathway and the PLP salvage pathway are intact and functional in *rus* mutants, the normal B6 vitamer levels are not surprising. Thus, another possible explanation for the vitB6-deficiency phenotypes of *rus* mutants is that there is a reduction in PLP availability. Since RUS1 and RUS2 are known as genetic and biochemical partners required for early seedling development, our expanded suppressor screens were conducted in both the *rus1* and the *rus2* mutant background. Our suppressor screens were effective, as a number of genetic suppressors were repeatedly and independently identified from both genetic backgrounds. We demonstrated in this study that PLP availability could indeed be restored in *rus1* and *rus2* mutants if specific changes are made in certain PLP-binding aminotransferases, which closely interacted with RUS1 and RUS2.

Our *in vivo* LC/ESI-MS/MS results showed that as much as 80% of the immunoprecipitated WT ASP2 proteins did not have PLP conjugation (Figure 5), suggesting that much of the ASP2 proteins exist as apoprotein monomers that are ready to take up PLP. Thus, there appeared to be more ASP2 protein in the plant than needed simply for aminotransferase activity. This is consistent with the lack of phenotypes seen in the various *sor* single mutants, and suggests that the seeming overabundance of these proteins may be due to them having a second function. We propose that the second function of aminotransferases is in the safe sequestration of PLP after its synthesis by the PLP *de novo* synthesis pathway. PLP homeostasis requires tight coordination among various biochemical processes including PLP synthesis, PLP sequestration/storage, PLP recovery and PLP usage. Our data suggest that *de novo* synthesized PLP is sequestered by specific aminotransferases, which are ubiquitous enzymes located in various cellular organelles. The dynamic binding and release of PLP from aminotransferases may serve as a mechanism to safely store PLP (Figure 9). Sequestration may both prevent PLP from forming inappropriate bonds with various biomolecules, and protect PLP itself from photo-degradation. The role of RUS1 and RUS2 in this working model is to interact with PLP-bound aminotransferases and facilitate the release of PLP for other cellular needs.

**Figure 9.**
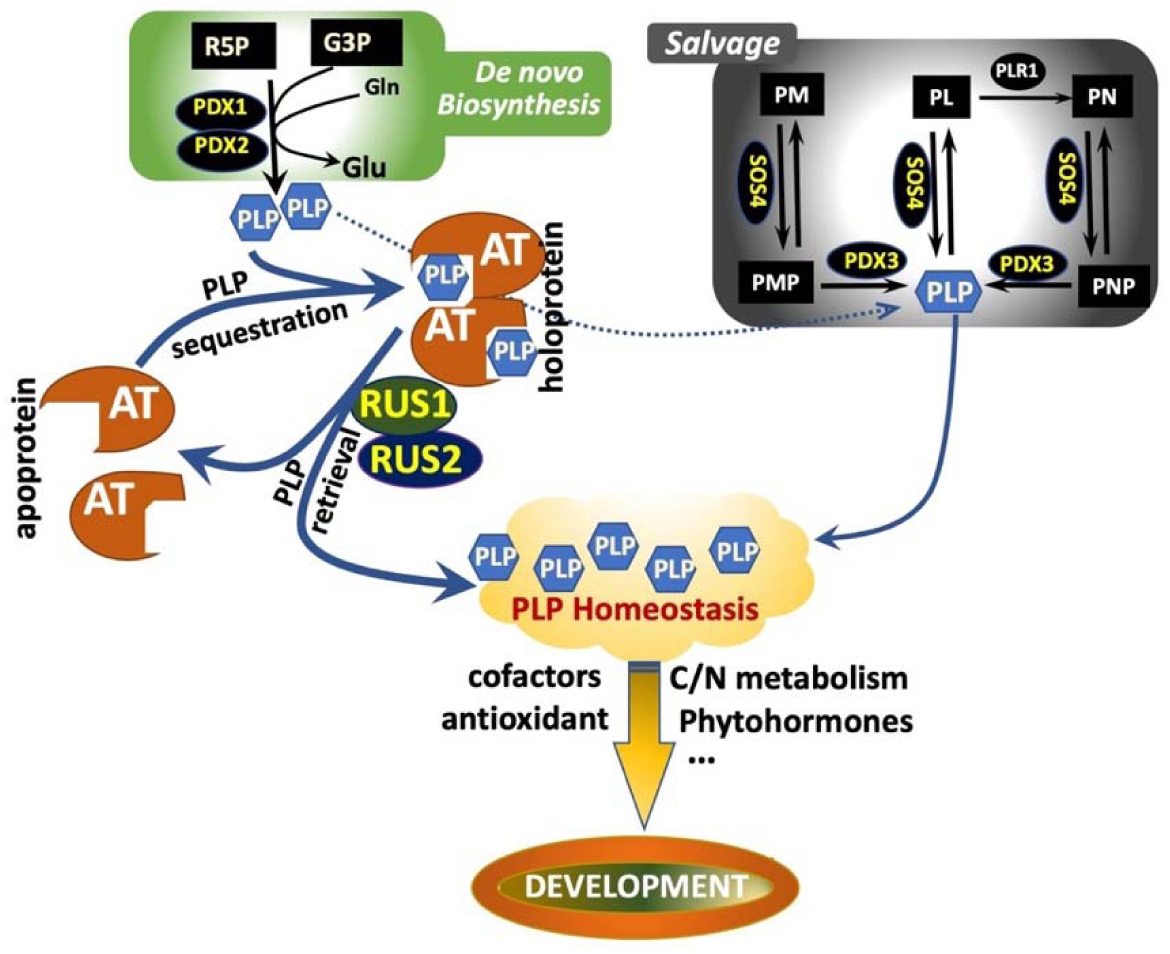
Model illustrating the roles of aminotransferases and RUS1/RUS2 in PLP homeostasis modulation. Maintaining PLP homeostasis is essential to plant development. PLP is an important vitamer used as a cofactor by a large number of enzymes and an antioxidant to cope with various stresses. PLP and PLP-dependent enzymes play critical roles in many biochemical processes including nitrogen metabolism and hormone synthesis. A healthy PLP homeostasis means PLP is continuously available while the free cellular PLP is kept low as accumulation of free PLP can be cytotoxic. We propose that PLP homeostasis is achieved by coordinated activities of both the de novo PLP biosynthesis pathway and the PLP salvage pathway. PLP synthesized de novo via the PDX1/PDX2 complex is sequestered by specific aminotransferases (AT). Upon binding to PLP, the AT apoprotein monomers dimerize to form holoprotein. RUS1 and RUS2 interact directly with ATs to make PLP available likely through regulating conformations of the PLP binding pockets in ATs. Biosynthesized PLP can also be potentially supplied to the PLP salvage pathway directly or indirectly. B6 vitamers generated during protein turnovers or gained by absorption from external sources can be interconverted via the PLP salvage pathway. The conversions of the non-phosphorylated forms (PM, PL, PN) to the phosphorylated forms (PMP, PLP, PNP) are catalyzed SOS4 (pyridoxal kinase). Both PNP and PMP can be converted to PLP by PDX3 (PMP/PNP oxidase).

## Materials and Methods

*Plant Material and Growth Conditions.* Arabidopsis plants were grown and handled as previously described (Hou et al., 2005). For Petri dish-grown seedlings, Arabidopsis seeds were surface-sterilized before cold-treated at 4°C for at least 48 h. Prepared seeds were plated on MS (Murashige and Skoog, 1962) growth medium in square plates (100x100x15 mm, Fisher Scientific) vertically held to allow seedlings grow on media surface so root morphology can be easily analyzed. For soil grown plants, cold-treated seeds were directly sowed in sterilized soil and kept in a growth chamber (Model AR-66L, Percival) with 16-h/8-h light/dark cycle at 22°C. Plants were imaged with a RT Slider camera using Spot software (Diagnostic Instruments, Inc.). High-resolution plant images were created on individual seedlings transferred to glass slides and images of different genotypes were put together using Adobe Photoshop (Adobe Systems, Inc.). Root-length measurements were performed using the ImageJ software (http://rsbweb.nih.gov/ij/). Mapping populations were created and selected as previously described (Leasure et al., 2011). Specific SSLP and CAPS markers were developed and used to map various *sor* alleles. Fine-mapping markers were used to map specific *sor* alleles to a 50-kb region that subsequent sequencing this region was performed to identify the mutation site.

Mutant Seeds and Genetic Crosses. Both rus1-2 and rus2-1 seeds were used to set up the EMS-based suppressor screens (Tong et al., 2008; Leasure et al., 2011). *pdx1.3-2*, *pdx3-3*, or *sos4-1*. The following mutant seeds were obtained from the Arabidopsis Biological Resource Center (ABRC, Ohio State University, Columbus, OH) (mutant alleles with their stock numbers): *pdx3-1* (SALK_060749), *pdx3-2* (SALK_149382C), *pdx3-3* (SALK_054167C) (Colinas et al., 2016), *sos4-1* (CS24930), *pdx1.3-2* (SALK_086418) (Ristilä et al., 2011). Homozygous double, triple or quadruple mutants were generated by multiple crosses and identified by genotyping with allele-specific PCR markers.

Protein Structure Analysis and Yeast Two-Hybrid. The Cn3D helper application (National Center for Biotechnology Information, NCBI) was used to place the positions of the substituted amino acid residues in the suppressor proteins based on the hGOT1 (PDB ID 30II) structure (Lee et al., 2018). Molecular Dynamics computer simulations and Pearson correlation coefficient (PCC) calculations were performed as previously described (Lee et al., 2018). Yeast two-hybrid assays and site-directed mutagenesis were performed as previously described (Leasure et al., 2009).

Protein Expression. The hGOT1 and mutant constructs were sub-cloned into the mammalian expression vector pcDNA3.1 (+), controlled by the CMV promoter and modified to possess a GFP at the C-terminus. The hGOT1-GFP and mutant hGOT1-GFP constructs were transfected into Chinese Hamster Ovary (CHO) (ExpiCHO-S) cells with an ExpiFectamine Transfection Kit (Gibco) following the manufacturer’s protocol. The transfected cell cultures were incubated on an orbital platform shaker (MaxQ 2000, Thermo Scientific) in a humidified incubator (MCO-20AIC, Sanyo) set to 32-37 °C and 5-8% CO2. After seven days, the cell cultures were removed from incubation and the cells were harvested for analysis. The cell pellets expressing the GOT1-GFP and its mutants were lysed with Cell Extraction Buffer (Invitrogen) mixed with a protease inhibitor cocktail (cOmplete Mini, Roche), and 1 mM phenylmethylsulfonyl fluoride. To fix covalently linked PLP cofactor, 5 mM sodium borohydride (Sigma) was dissolved into the lysis solution. Cell lysis proceeded on ice for 30 minutes with brief vortexing at 10 minutes intervals. The cell lysate was centrifuged at 20800 x g for 15 minutes, and the resulting supernatant was collected for SDS-PAGE or immunoprecipitation experiments.

*Immunoprecipitation*. Pulldowns of the GFP-fusion proteins were performed by incubating tissue homogenate (7-day-old seedlings) or cell lysate (CHO cell lysate) with agarose beads that conjugated with GFP antibodies (GFP Nano-Trap, Chromotek). After incubations of 4-24 hours with gentle inverting, the GFP agarose beads were collected in a Mobicol “F “column (MoBiTec) and washed with PBS at 4 °C. Single column volumes of a 44 mM sodium acetate, pH 3.5 solution was applied to the column and the eluate was collected into fractions. Each fraction was neutralized with HEPES to a final pH of 7.0. The absorbance at 280 nm was determined for each fraction with a Nanodrop 2000c (Thermo Scientific).

### Protein SDS-PAGE and Trypsin Digestion

Protein samples were fractionated on 10% or 4-12% acrylamide Bis-Tris gels (NuPAGE, Novex). Extracted proteins were prepared in reducing SDS buffer with 50 mM DTT and incubated at 70 °C for 10 minutes. Samples were loaded in the electrophoresis chamber (XCell SureLock, Thermo Scientific) with 1x MES-SDS buffer and ran at 120 V for 50 min. Gels were washed with distilled water, and stained with Coomassie G-250 (SimplyBlue, Thermo Scientific) for 3 hours. Gels were de-stained overnight with distilled water. Aqueous protein fractions or isolated protein bands were excised and de-stained with 50% acetonitrile, 25 mM NH_4_HCO_3_ solution. Protein bands were reduced with 50 mM DTT in a 50 mM NH_4_HCO_3_ wetting buffer at 55 °C for 25 minutes, washed, and alkylated in 60 mM IAM in wetting buffer without light at 37 C° for 25 minutes. Protein gel bands were dried without heat in a vacuum concentrator (SpeedVac Plus, Savant). The protein was resuspended in wetting buffer with 1:20 trypsin protease (Pierce) to protein (w/w) at 37 °C overnight.

*Protein Identifications by LC/ESI-MS/MS Analysis*. Tryptic peptides derived from trypsin-digested samples were dissolved with 0.1% formic acid/water, and subsequently analyzed by liquid chromatography/electrospray ionization-tandem mass spectrometry (LC/ESI-MS/MS) using a hybrid quadrupole orbitrap mass spectrometer (QExactive, Thermo Scientific) coupled with a LC system (Dionex Ultimate 3000, Thermo Scientific). LC/ESI-MS/MS analyses were conducted using a reverse phase C18 column (75μm x 130mm). The mobile phases for the reverse phase chromatography were (mobile phase A) 0.1% HCOOH/water and (mobile phase B) 0.1% HCOOH in acetonitrile. A 90min multiple steps gradient program was used for LC separation (holding at 5% B for the first 10min, increasing to 12.5% B in 5min, ramping up to 35% B in the next 40min, followed by increasing from 35% to 80% B in the next 10 min, holding at 80% B for 7 min, and back to 5% B during the final 10 min). The ESI-MS/MS data acquisition was set to collect ion signals from the eluted peptides using an automatic, data-dependent scan procedure in which a cyclical series of 11 different mass scan modes was performed (a full MS scan followed by 10 MS/MS scans for the top ten most abundant ions). The resolution of QExactive was set at 70,000 for the full MS mode, and 17500 for the MS/MS mode, respectively. The full mass range was set at a scan range from m/z 400 to 1600 and the dynamic exclusion duration for MS/MS acquisition was set at 10s. The protein identification algorithm Sequest in Proteome Discoverer program was used to identify peptides from the resulting MS/MS spectra by searching against the protein database extracted from SwissProt (v57.14; 2010 February) including the sequences of the known mutants. Searching parameters were set as parent/fragment ion tolerances of 20 ppm/0.1Da. Carbamidomethyl modification of Cys was set as a fixed modification. The reduced form of Pyridoxal Phosphate H2 at Lys were set as variable modifications. The maximum missed cleavage of trypsin was set as 2. Scaffold (Proteome Software, v4.2.1) was used to merge and summarize the LC/MS/MS data. The protein identification was set to have a minimum of two peptides with a false positive rate of 1% for the protein identification based on the decoy data base searching.

*Bimolecular fluorescence complementation (BiFC) assay*. Full-length cDNA coding sequences (without the stop codon) of both RUS1 and ASP2 were PCR amplified using gene-specific primers. The amplified full-length cDNAs were cloned into pENTR/SD/D-TOPO vectors (Invitrogen), and then subcloned between the 35S promoter and YFP coding sequence in the destination plasmid pEarleyGate 101, using Gateway recombination cloning reaction (Thermo Fisher). The resulted fusion constructs were obtained and confirmed by sequencing before transformed into agrobacterium strain GV3101. Agrobacterial cells carrying the various constructs (CCFP-RUS1, NYFP-RUS2, NYFP-ASP2) were cultured. Transformed agrobacterial cells are infiltrated to tobacco leaf tissues according to the previously reported procedures (Schweiger and Schwenkert, 2014). BiFC GFP signals were detected by a Zeiss LSM 710 Confocal microscope with a Plan-Apochromat 20x/0.8 objective lens. A blue laser (488 nm) was used for excitation. Channel 1 (495-550 nm) was used for GFP detection and Channel 2 (650-705 nm) was used for chlorophyll autofluorescence detection. Images processed by both ZEN software (Zeiss) and FIJI were arranged by Photoshop.

## Acknowledgements

This work was partly supported by NIGMS (National Institute of General Medical Sciences) Award SC1GM095462, NIH grant S06 GM52588, and NIH MBRS-RISE R25-GM059298.

## Author Contributions

H.T., C.D.L. and Z.-H.H. designed the experiments; H.T. performed most of the experiments; R.Y. performed the mass spectrometry analyses. J.L. and A.G. performed the Molecular Dynamics analysis. N.O performed the genetic tests for SOS4 and PDX3. D.T. performed experiments with the hGOT. Y.S. and S.Z. performed the immunoprecipitation experiments and analyzed the MS results. D.B. performed the GUS analyses. Y.T., E.M.D., S.P., C.I., J.-T.C. T.M., N.P., X.P., and K.T. performed the genetic tests for various suppressors and analyzed phenotypes. H.T., C.D,L. and Z.-H. H. analyzed the data. C.D,L. and Z.-H. H. wrote the article.

